# Altering the bilayer motif in ERBB2 HER2 TMD and in ErbB HER TMD dimer causing in vitro and in vivo tumor suppression

**DOI:** 10.1101/2020.03.05.978080

**Authors:** Laszlo David Menyhert, Miguel Tejeda

## Abstract

Human ERBB2 is a transmembrane signaling tyrosine kinase receptor, which seems an ideal target of human WNT16B, the secreted growth factor possibly causes transmembrane domain (TMD) mutations. There is a strong relationship between the chemical nature of the TMD mutations and the potency with which they activate HER2. *In silico*, we modeled the possible docking conformation of human WNT16B and human ERBB2 TMD homodimer, resulted a mutant complex. The ribbon structure, the C-terminal and N-terminal and GG4-like motif structures are similar in HER2 TMD and HER TMD, we modeled WNT16B’s possible docking conformation to the HER1 TMD (ErbB), also resulted a mutant complex. If there is a strong relationship between TMD mutations improving the active dimer interface or stabilizing an activated conformation and the potency with which they activate HER2 (and possibly also HER), than the TMD dimerization part seems ideal reagent-target. The agent we tested – the 4-(Furan-2-yl)hepta-1,6-dien-4-ol (AKOS004122375) – has very good connectivity attributes by its several rotatable bonds, and according to the *in silico* inspection of close residues intermolecular bonds, and the ligand docking, it can straight connect to human ERBB2 TMD (HER2), and to the ErbB TMD (HER1) dimer bilayer motif as well. *In silico*, we also tested the agent ligand’s docking into the residues of human WNT16B and human ERBB2 TMD (HER2) mutant complex, and human WNT16B and human ErbB TMD (HER1) mutant complex. We tested the agent ligand *in vitro* and *in vivo* in several tumor models, highlighting that targeting the EGFR’s TMD with an agent not only reduces treatment-induced metastasis, but radically decreases the tumor growth as well. Because of the analogous structure of HER2 TMD and HER TMD, this dimerization motif-targeting can also be successful in HER and HER2 EGFR signaling. *In vitro*, we reached 80-94% proliferation percentage in different tumor models, *in vivo* we reached 35-61% tumor suppression in different tumor models, the metastasis inhibition effect of the compound was 82-87% in different tumor models.

## Introduction

The development of cancer cells is a process, and when the normal regulation of cell division goes bad, long series of typical mutations lead to cancer development. The aging of cells is a biological mechanism, which is supervised by the p53 / p21 (WAF1) stimulus path, and some factors are overexpressed in fibroblasts in those cells, that undergo repeated phases of aging. Both p53 and the growth-factor protein require the development of replicative cellular senescence, and also the growth-factor protein regulates phosphoinositide 3-kinase (PI3K) / AKT pathway activation, which is linked via the epidermal growth factor receptor (William, 2010; Joong & Jun, 2014). In a relevant number of cases, the cancer cells become resistant to anticancer agents or toxins during chemotherapy. WNT16B protein has the immoderate information of generating strong metastasis (Weinstein & Buolamwini, 2000; Schwartsmann *et al*, 2002, Kapuvári *et al*, 2010). The relevant publications marked, that this growth factor protein’s level can be up to thirty times more in the body of a patient being treated with chemotherapy, than in a healthy one (Sun Y *et al*, 2012; Muy-Teck *et al*, 2007).

Deregulated HER2 is a target of many approved cancer drugs, structural modeling and analysis showed that the TMD/JMD mutations function by improving the active dimer interface or stabilizing an activating conformation (Pahuja *et al*, 2018). During signal transduction across the plasma membrane, ErbB receptors are involved in lateral homodimerization and heterodimerization with proper assembly of their extracellular single-span transmembrane (TM) and cytoplasmic domains (Mineev *et al*, 2014). In humans, more than 70% of ErbB2-positive sporadic breast cancers harbor p53 mutations, which correlate with poor prognosis, and in addition, the extremely high incidence of ErbB2-positive breast cancer in women with p53 germ-line mutations (Li-Fraumeni Syndrome) suggests the key role of mutant p53 specifically in ErbB2-mediated mammary tumorigenesis (Yallowitz *et al*, 2015).

A possible docking protein of the HER2 TMD homodimer receptor site is WNT16.This protein’s expression in nearby normal cells is responsible for the development of chemotherapy resistance, the chemotherapeutic patient’s body could contain thirty times more WNT16B than normally. These proteins are glycosylated, and associated with the cell surface or transmembrane matrix, recognize cell surface receptors. WNT16B contains two transcript variants diverging at the 5’ termini, these two variants are proposed to be the products of separate promoters and not to be splice variants from a single promoter. They are differentially expressed in normal tissues, one of which (variant 2) is expressed at significant levels only in the pancreas, whereas another one (variant 1) is expressed more ubiquitously with highest levels in adult kidney, placenta, brain, heart, and spleen. WNT16B expression is regulated by nuclear factor of κ light polypeptide gene enhancer in B cells 1 (NF-κB) after DNA damage, as can occur in normal cells during radiation or chemotherapy. Subsequently, WNT16B signals in a paracrine manner to activate the Wnt expression program in tumor cells.

**Figure 1.**
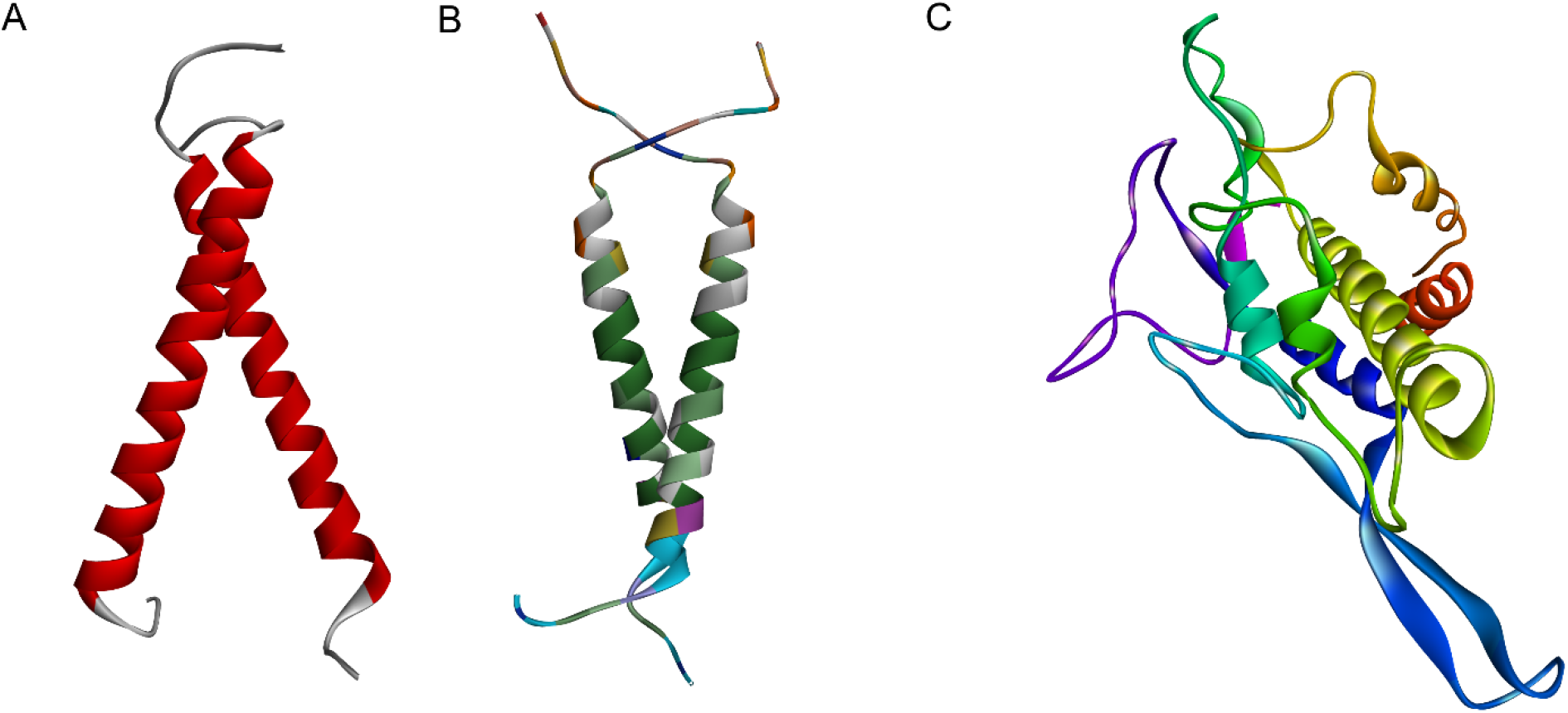
(A) ERBB2 HER2 TMD homodimer (PDB: 2JWA) (B) ErbB HER TMD homodimer in micelles (PDB: 2M0B) human transmembrane tyrosine kinase receptor segments in active form (C) human WNT16B growth factor protein

## Results and Discussion

Human ERBB2 is a transmembrane signaling tyrosine kinase receptor of EGF, which seems a possible target of human WNT16B (Zhang *et al*, 2015), the ErbB family contains four plasma membrane-bound receptor tyrosine kinases. ERBB2tm dimer structure (HER2 TMD) explains the biochemical and oncogenic properties of human ERBB2 receptor and provides a basis controlling its kinase activity, which is critical in many disease states, proper lateral dimerization of the transmembrane domains of receptor tyrosine kinases is required for biochemical signal transduction across the plasma membrane (Bocharov *et al*, 2008). Additionally, specific protein-protein and protein-lipid interactions of singlespan HER transmembrane domains (TMDs) are important for proper receptor activation and mechanism(s) that reduce or enhance such interactions (e.g., by means of mutations) and can affect downstream activity independently of KD mutations (Ou *et al*, 2017). Relevant publications observed a striking relationship between the chemical nature of the TMD mutations and the potency with which they activate HER2 (Pahuja *et al*, 2018). Another relevant publication uses the HER2 TMD homodimer model (PDB: 2JWA) as HER TMD JM-A homodimer (Jura N *et al*, 2009): the transmembrane part of the two pathways can be similar, ribbon structures of the HER TMD dimers formed via the alternative C-terminal and N-terminal and GG4-like motifs, just like in HER2 TMD homodimer (Bocharov *et al*, 2016; PBD: 2M0B). So it is possible, that the TMD mutations both affecting HER (Cymer & Schneider, 2010) and HER2 through the way is reviewed in this study.

By using expression profiling techniques, it was managed to search for secreted factors, that were overexpressed in fibroblasts undergoing replicative senescence (Smolich *et al*, 1993). WNT16B is overexpressed in cells undergoing stress-induced premature senescence and oncogene-induced senescence in both MRC5 cell line and the *in vivo* murine model of K-Ras(V12)-induced senescence, his secreted factor seemed a probable docking protein of HER2 TMD (Binet *et al*, 2009). By small interfering RNA experiments, it was observed, that both p53 and WNT16B are necessary for the onset of replicative senescence, and WNT16B expression is required for the full transcriptional activation of p21 (WAF1). Moreover, WNT16B regulates activation of the phosphoinositide 3-kinase (PI3K)/AKT pathway, so overall, WNT16B is a new marker of senescence that regulates p53 activity and the PI3K/AKT pathway and is necessary for the onset of replicative senescence (Binet *et al*, 2009). Furthermore, if HER TMD homodimer has the same structure as HER2 TMD homodimer (Jura N *et al*, 2009), than WNT16B could affect HER and HER2 both.

**Figure 2.**
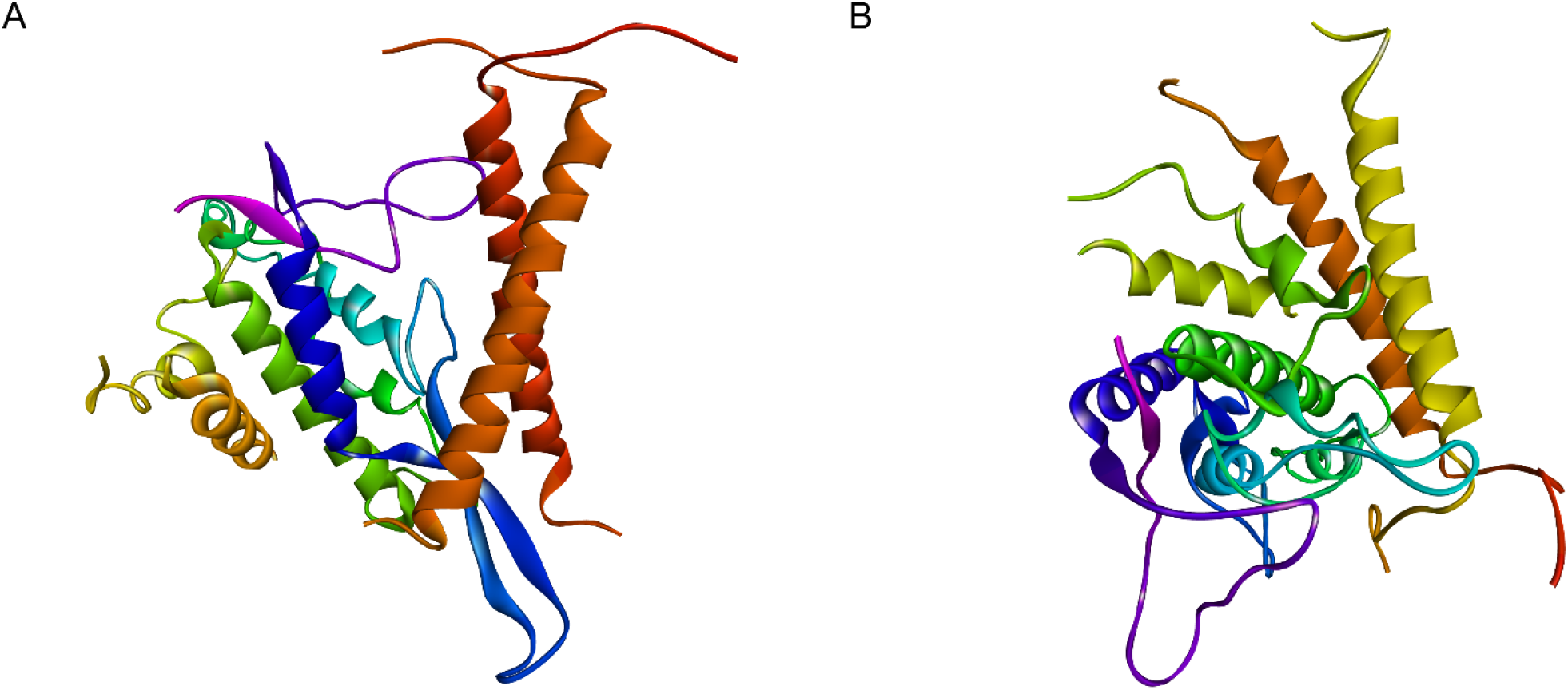
Possible conformations of human WNT16B is docking into the HER2 TMD homodimer (A) and HER TMD homodimer (B) receptor sites

*In silico*, after reviewing the close residues intermolecular bonds of HER2 TMD homodimer (PDB:2JWA), and human WNT16B, we experienced, that A:ARG77, A:GLN79, A:GLN80, A:GLN80, A:ILE82, A:ARG83, A:ARG83, A:LYS84, B:PHE171, B:ILE175, B:LYS176 parts of HER2 transmembrane domain can bind with conventional hydrogen intermolecular bonds to WNT16B homo sapiens as H-donors, A:LYS, A:LEU, A:THR and the A:VAL parts of human WNT16B also behave as conventional hydrogen H-donors, striking indicates they mutate very probable. We also reviewed, the carbon hydrogen and other intermolecular bonds between HER2 TMD and human WNT16B, and the possible intermolecular bonds between the complexes show a very strong binding, docking potential.

*In silico*, we also reviewed the close residues intermolecular bonds of HER TMD homodimer (PDB: 2M0B), and human WNT16B, we experienced, that the A:ARG, A:LEU, B:ARG, B:ILE, B:LYS parts of HER1 transmembrane domain can bind to human WNT16B as H-donors with intermolecular conventional hydrogen bonds, human WNT16B’s A:THR, A:GLU, A:ASN parts behave as H-donors in conventional hydrogen bonds. We also reviewed the electrostatic and hydrophobic intermolecular bonds as well, the results indicate they mutate very probably.

**Figure 3.**
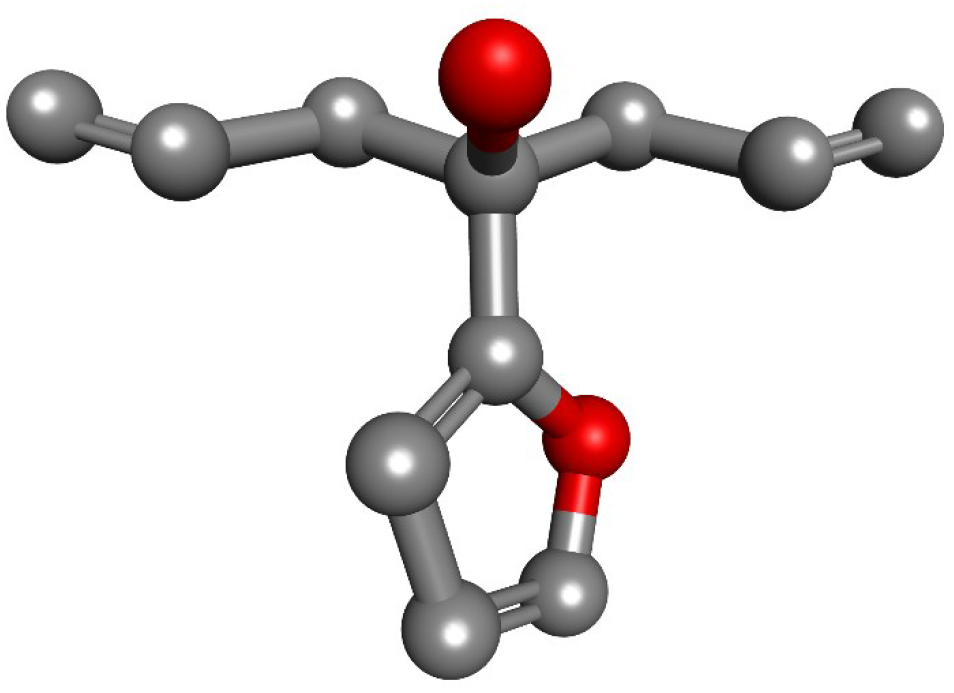
The compound 4-(Furan-2-yl)hepta-1,6-dien-4-ol (AKOS004122375)

The compound AKOS004122375 (InChIKey: YTNUPDNFKZGGB-UHFFFAOYSA-N, PubChem SID: 108669579) is a form of 4-(furan-2-yl)hepta-1,6-dien-4-ol (alpha,alpha-diallylfuran-2-methanol, PubChem CID: 11206108). The molecular weight is 178.23 g/mol, the compound has 1 hydrogen bond donor, and 2 hydrogen bond acceptor sites, with 5 rotatable bonds, the formal charge is 0. The differences between the sub-ligands of the 4-(furan-2-yl)hepta-1,6-dien-4-ol only affect the free transform properties of the rotatable bonds. The ERBB2 transmembrane domain (HER2 TMD) has a homodimer and a hetero-dimer form (Bocharov *et al*, 2008), it can be inactive or active conformed, in this study we only reviewed the active form of the homodimer. If the HER TMD homodimer has the same structure as HER2 TMD homodimer (Jura N *et al*, 2009), than probably the same interaction can be observed at the dimerization-bilayer motif of HER TMD.

When focusing on the intermolecular bonds of the close residues of AKOS004122375 and HER2 TMD homodimer (PDB:2JWA), it can be seen that the 4-(furan-2-yl)hepta-1,6-dien-4-ol:O1 atom bound with conventional hydrogen bond to HER2 TMD A:VAL64, the distance is 3.02086,. Furthermore, the compound also bound with three carbon hydrogen, and two Pi-alkyl bonds to HER2 TMD. When reviewing the close residues intermolecular bonds between the agent compound AKOS004122375 and human WNT16B, the ligand bound with four Pi-alkyl bonds to HER TMD A:LEU and A:ALA parts, with distance of between 4.1 and 5.1.

**Figure 4.**
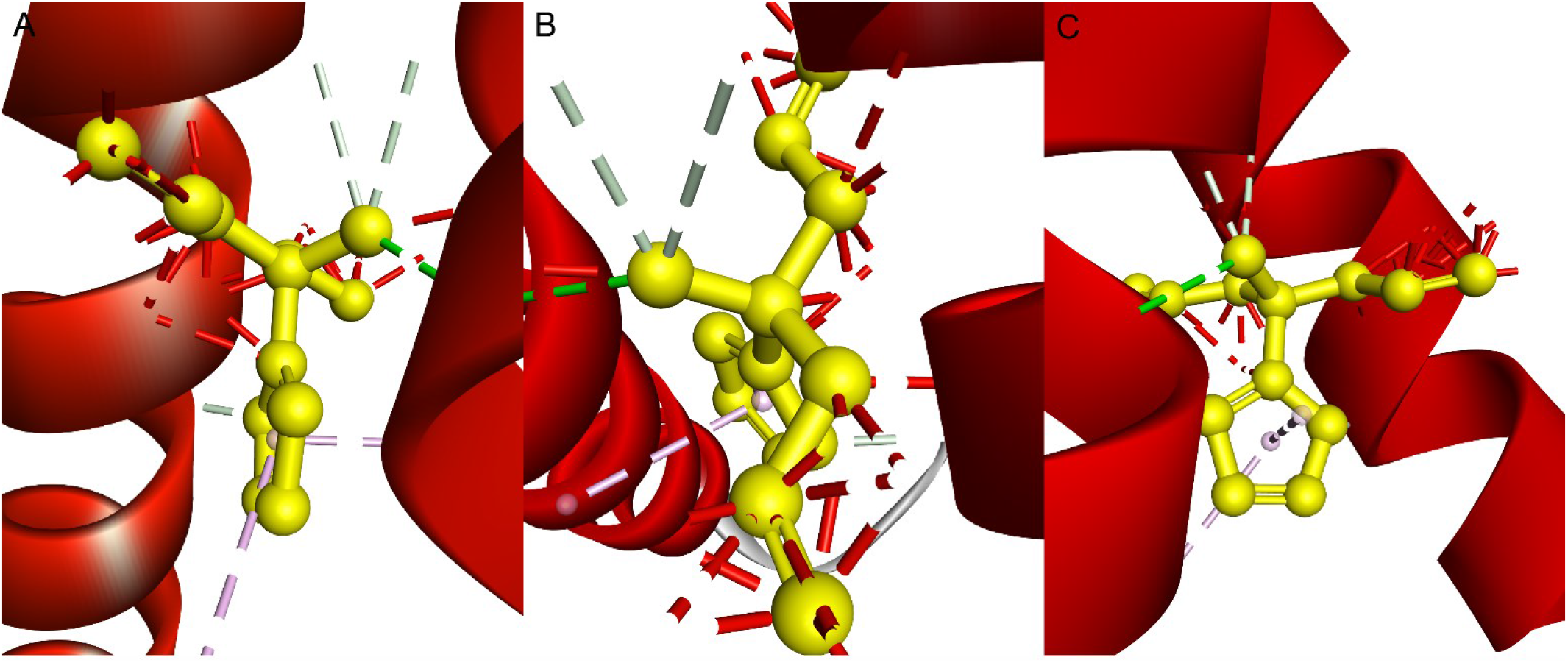
Possible (A) intermolecular bonds of the close residues of 4-(Furan-2-yl)hepta-1,6-dien-4-ol (AKOS004122375) and HER2 TMD homodimer, (B) bonds of AKOS004122375 is docking into the HER2 TMD homodimer and human WNT16B mutant complex, (C) bonds of AKOS004122375 is docking into the HER2 TMD homodimer

We tested *in silico* the possible results of AKOS004122375 ligand docking into the HER2 TMD homodimer (PDB:2JWA), and human WNT16B mutant complex, our results show that the ligand bound the same way when docked into only the HER2 TMD. Importantly, HER2 inhibiting antibodies and small molecules can block the activity of HER2 transmembrane domain and juxtamembrane domain mutants (Pahuja *et al*, 2018), our agent ligand acts exactly like a kind of inhibitor. The exact mechanism will be investigated later, but it is possible, that the agent compound alters the partial and surface charge of the helix (especially in the dimerization-bilayer motif), which causes probable malfunction in bad signaling mutation pathways.

**Figure 5.**
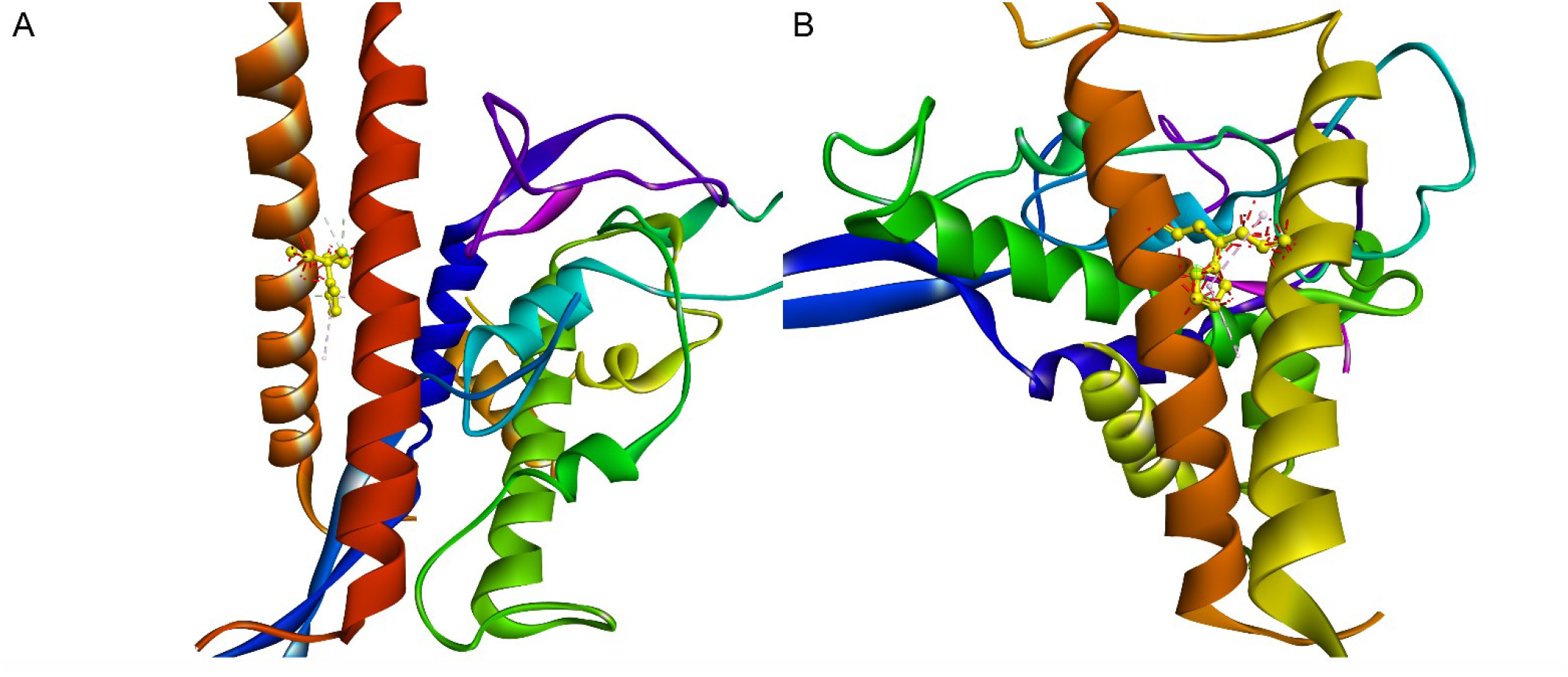
Possible conformations of 4-(Furan-2-yl)hepta-1,6-dien-4-ol (AKOS004122375) is docking into the HER2 TMD homodimer and human WNT16B mutant complex (A) and into the HER TMD homodimer and human WNT16B mutant complex (B)

The reagent compound (AKOS004122375) behaves H-acceptor, when connected to human HER2 TMD homodimer (PDB:2JWA) A:VAL64 with conventional hydrogen bond (distance: 3.02086, angle xda: 112.19, angle day: 125.117). The HER2 TMD A:GLY60, B:GLY160 and B:VAL:164 behaving H-donor when bound to 4-(furan-2-yl)hepta-1,6-dien-4-ol (AKOS004122375) with carbon hydrogen bonds (3 bonds with 1.87114-3.043 distances). Furthermore, there are two Pi-alkyl bonds between the compound and HER2 TMD’s A:VAL64 and A:LEU67 parts, with above 3.2 in distance. The compound AKOS004122375 bound with Pi-alkyl intermolecular bonds to A:ALA661, B:LEU657, B:LEU658 parts of HER TMD, with distance of between 4.1 to 5.1.

**Figure 6.**
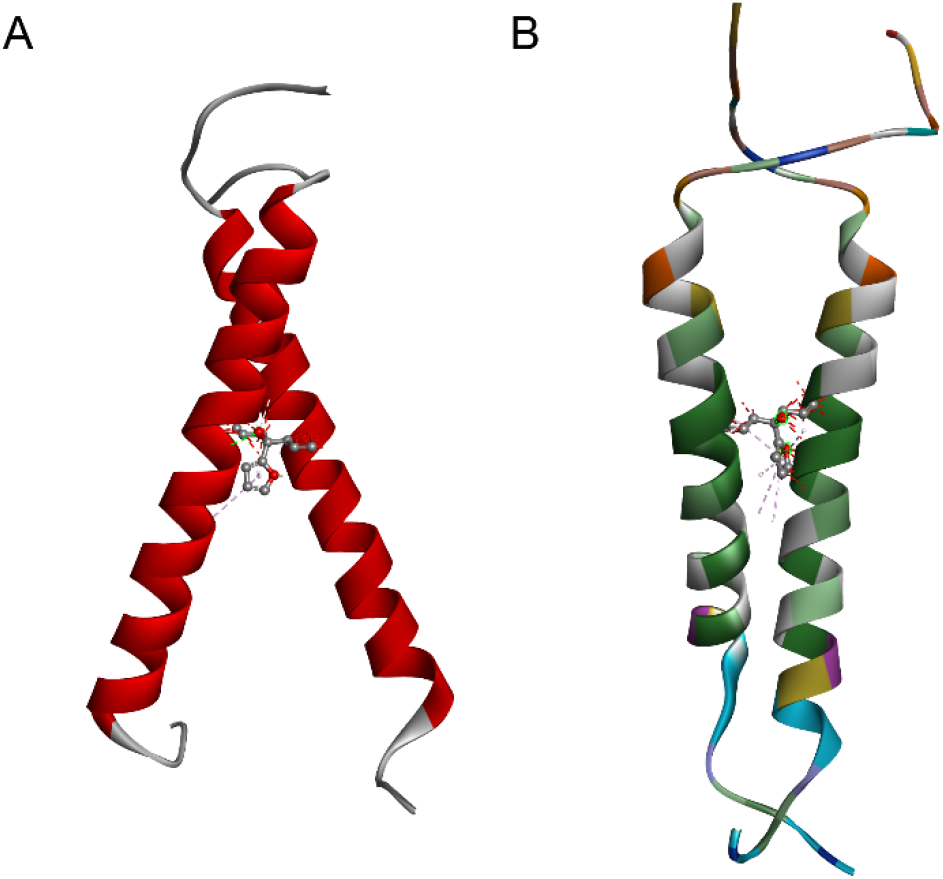
Possible conformations of 4-(Furan-2-yl)hepta-1,6-dien-4-ol (AKOS004122375) is docking into the HER2 TMD (A) and into the HER TMD (B)

Generally, we can assert, that – according to our *in silico* experiments – human WNT16B is very likely to dock into the HER and HER2 TMD homodimer, and this could cause an act on the biochemical signal transduction across the plasma membrane. This could affect the p51, p53, PI3K pathways (William, 2010; Joong & Jun, 2014), and lead to the confusion of normal cell functions, possibly causes the repeated phases of cell aging. Our theory, a simple ligand could change the crucial binding structure of the secreted growth factor and the tyrosine-kinase receptor, through strong connection to the active form of HER and HER2 TMD dimerization motif, is demonstrated in our *in vitro* and *in vivo* experiments. We suggest a key role for ErbB ErbB2-mediated mutant p53 in treatment-induced metastasis, and as well in tumor growth.

*In vitro* we tested the antiproliferative efficacy of 4-(Furan-2-yl)hepta-1,6-dien-4-ol (AKOS004122375) on different human tumor cells: *in vitro*, the agent inhibited the proliferation of breast carcinoma cells MCF-7 and that of MDA-MD-231. Through MTT assays, the inhibitory concentration 50 (IC50) was determined at 72 h of exposure. Of MCF-7 cells the IC50 was 24O±l2.6 nM, and for MDA-MD-231 cells the IC50 was 180±20.6 nM. Of HT168-M1 cells the IC50 was 210+10.0 and for B16 cells the IC 50 was 175+19.2 nM. the antiproliferative effect of the new agent was observed in both cell lines at the exposure of 24 h. In 1.2 concentration, the proliferation percentage was in B16 cells 80%, in MCF-7 cells 87%, in MDA-MB 231 cells 94%, in HT168-M1 cells 89%.

**Figure 7.**
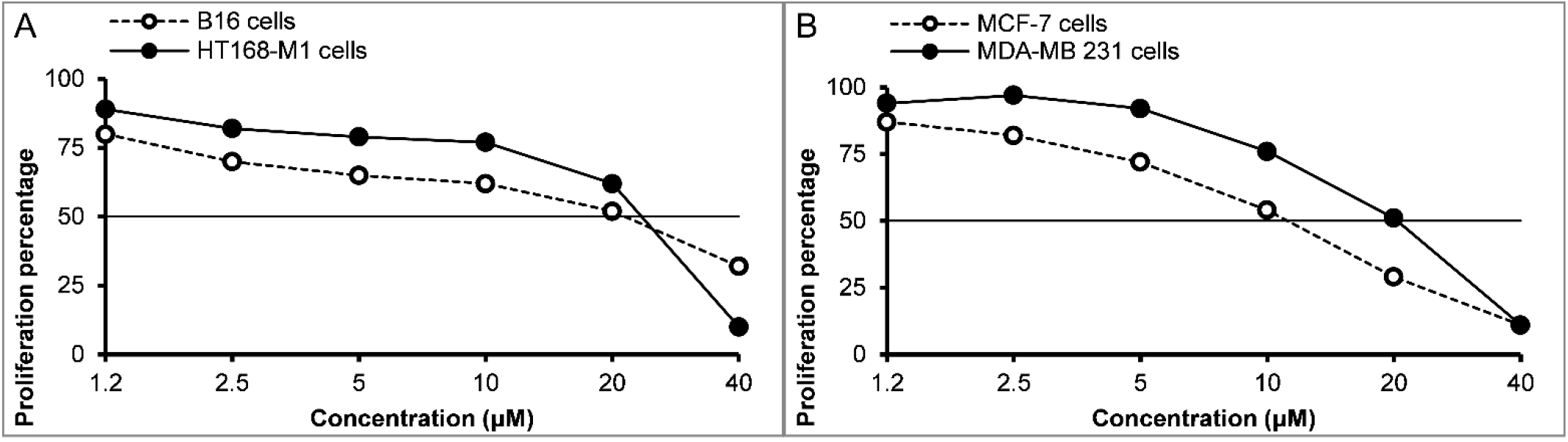
*In vitro* results of the proliferation effect of 4-(Furan-2-yl)hepta-1,6-dien-4-ol (AKOS004122375) on (A) B16 and HT168-M1 tumor cells, and on (B) MCF-7 and MDA-MB 231 cells

The toxic effect of 4-(Furan-2-yl)hepta-1,6-dien-4-ol (AKOS004122375) was studied in healthy mice on their weights, in different organs (lung, heart, liver, spleen, kidneys), with the application of 3 different doses. The new antitumor agent was not lethal for the experimental animals, only slightly decreased the weights of mice. The results demonstrated that the molecule was not toxic. The cytostatic and antiproliferative effect of 4-(Furan-2-yl)hepta-1,6-dien-4-ol (AKOS004122375) was also determined by MTT-assay (3-(4,5-dimethylthiazol-2-yl)-2,5 diphenyltetrazolium bromide), which is based on the conversion of MTT to aformazan derivative only by the living cells (Leurs *et al*, 2011).

The *in vivo* antitumor inhibitory effect of 4-(Furan-2-yl)hepta-1,6-dien-4-ol (AKOS004122375) was investigated in M1 human leucemia, M1 human melanoma, B16 F10 human melanoma, S-180 sarcoma, colon-26 adenocarcinoma, MXT mammary carcinoma, and B-16 melanoma tumor models, we reached the best results with the smallest dose. The treatment of AKOS004122375 4-(Furan-2-yl)hepta-1,6-dien-4-ol was carried out with 3 different doses (80mg/kg, 40mg/kg and 10mg/kg) via injections and applied twice a day for 2 weeks (Tejeda M *et al*, 1999). It caused an average 50%-70% decrease in the tumor volume in mice. The best results were shown at 10mg/kg dose (the smallest dosage). We’d also monitored the metastasis effects, which were low.

Figure 8. and 9. presents the *in vivo* antitumor activity of AKOS004122375, at three doses (10, 20, 40mg/kg), with subcutaneous injection and a 14xqd treatment schedule. Treatment with the agent started when the tumor had developed and became measurable. Mice were randomized before the beginning of treatments. The therapeutic effect of the agent was compared and expressed in terms of tumor growth inhibition and life span. Figure 8. (A) demonstrates the effect of AKOS004122375 on Colon-26 adenocarcinoma tumor model. On the basis of tumor growth curves and mean tumor volumes significant (35%) growth inhibitory efficacy of tumor development was observed with 10mg/kg, s.c. dose. When we applied AKOS004122375 at 20mg/kg and 40mg/kg ca. 21% tumor inhibitory effect was achieved. In our experiments we studied the antitumor effect of the compound on M1 human leukemia tumor model with three doses. Figure 8. (B) showing the doses (10mg/kg, 20mg/Kg and 40mg/Kg) resulted in 47% and 34% and 14% tumor inhibitory effects. In Figure 8. (C) we studied the therapeutic efficacy of the compound on MXT mammary carcinoma tumor model. When AKOS004122375 was administered at 10mg/kg, s.c. dose the agent induced a significant (42%) tumor growth-inhibitory effect in MXT mammary carcinoma tumor development. in Figure 8. (D) we investigated the inhibitory efficacy of the ligand on S-180 sarcoma tumor model following doses 10mg/kg, 20mg/kg and 40mg/kg. The tumor inhibitory activity of the agent at the end of the 14 days treatment period was 47% (10mg/kg), 43% (20mg/kg) and 30% (40mg/kg).

**Figure 8.**
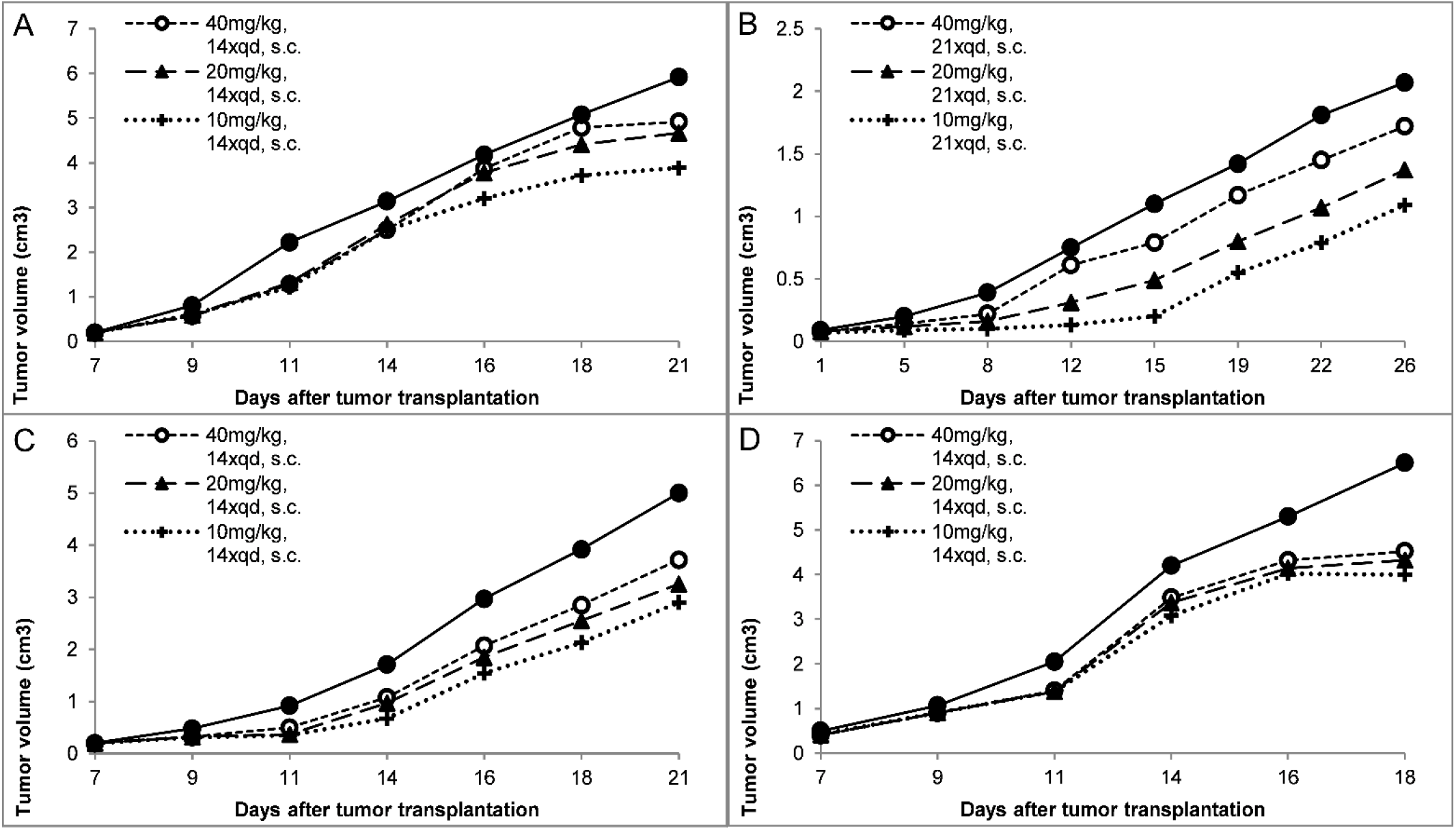
*In vivo* results of 4-(Furan-2-yl)hepta-1,6-dien-4-ol (AKOS004122375) tumor suppressor effects on (A) Colon-26 adenocarcinoma tumor model, on (B) M1 human leucemia tumor model, on (C) MXT mammary carcinoma tumor model, and on (D) S-180 sarcoma tumor model

The antitumor effect of AKOS004122375 on B16F10 melanoma tumor model was investigated with three doses: 10, 20 and 40 mg/kg with s.c. treatments. On the basis of tumor growth curves, a significant antitumor activity of the agent was observed. The antitumor activity of AKOS004122375 following s.c. treatments for 2 weeks was 61% (10mg/kg), 31% (20mg/kg) and 18% (40mg/kg) (Figure 9.).

**Figure 9.**
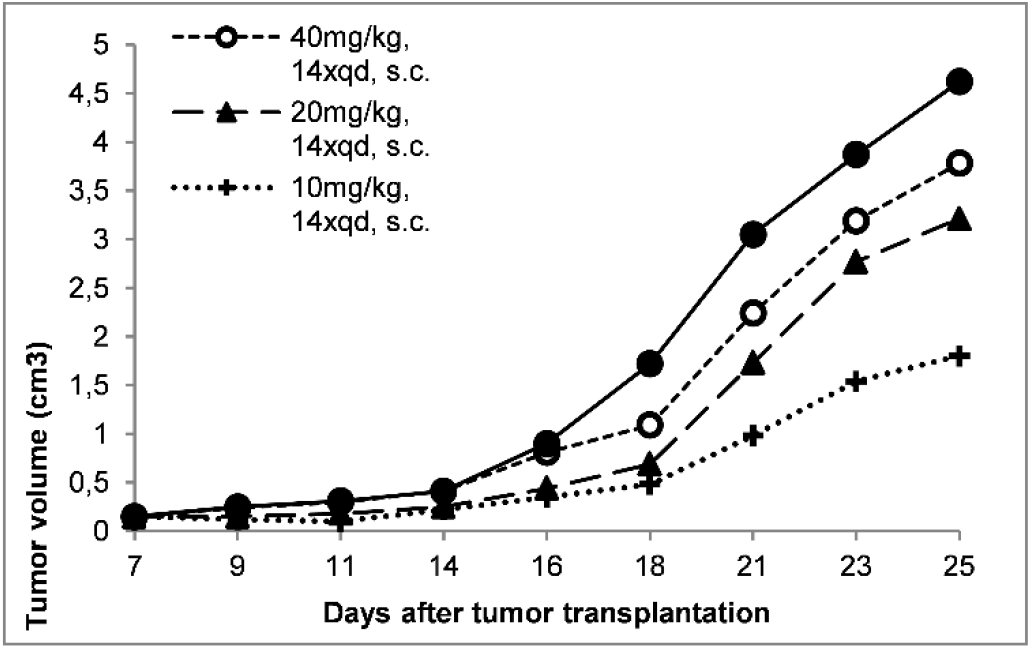
*In vivo* results of 4-(Furan-2-yl)hepta-1,6-dien-4-ol (AKOS004122375) tumor suppressor effects on B16F10 melanoma tumor model

The metastasis inhibition effect of the compound, we tested the liver metastasis formation of the M1 human melanoma tumor (mean± SEM) and on lung metastasis formation of the B16F10 melanoma tumor (mean± SEM with application of three different doses (10, 20 and 40mg/kg), with i.p. injection and 18xpd treatment schedule. The results demonstrated that all three doses significantly inhibited the number of metastases in comparison with the untreated control group. In the case of the M1 melanoma tumor 82% (10mg/kg), 63% (20mg/kg) and 44% (40mg/kg) inhibition was observed. In the case of B16F10 tumor 87% (10mg/kg), 79% (20mg/kg) and 72% (40mg/kg) lung metastases inhibition we have achieved (Figure 10).

**Figure 10.**
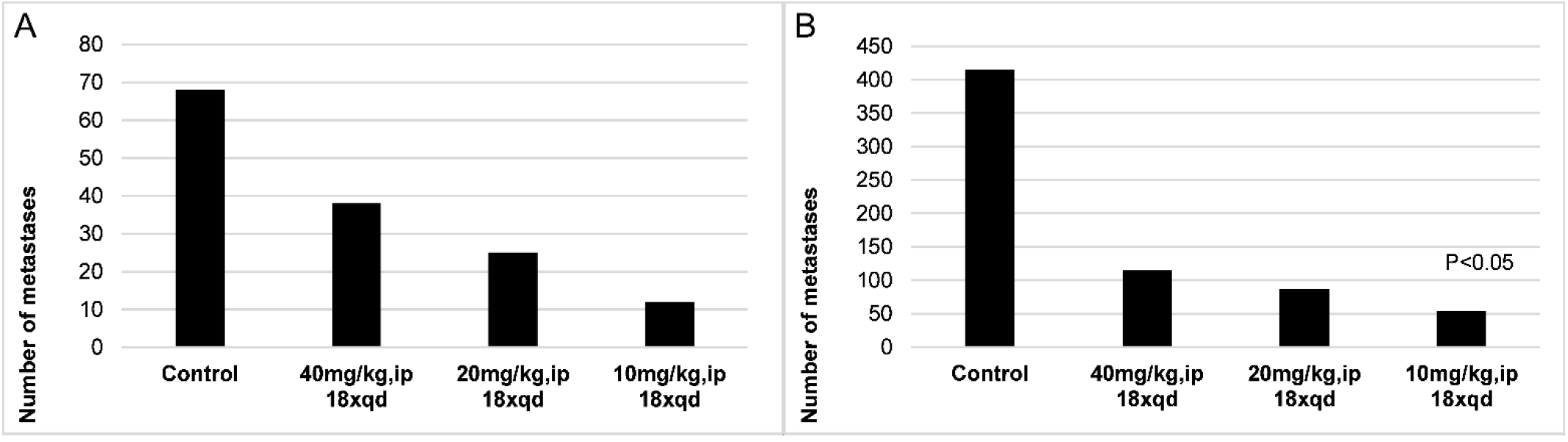
*In vivo* results of 4-(Furan-2-yl)hepta-1,6-dien-4-ol (AKOS004122375) metastasis suppressor effects on (A) liver metastasis formation of the M1 human melanoma tumor (mean± SEM) and (B) on lung metastasis formation of the B16F10 melanoma tumor (mean± SEM)

## Materials and Methods

The common feature of molecular modeling techniques is the atomistic level description of the molecular systems. This may include treating atoms as the smallest individual unit (the molecular mechanics approach), or explicitly modeling electrons of each atom (the quantum chemistry approach). Molecular modeling and synthesis encompass all theoretical methods and computational techniques used to model or mimic the behavior of molecules. Techniques are used in the fields of computational chemistry, drug design, computational biology and materials science for studying molecular systems ranging from small chemical systems to large biological molecules and material assemblies. During the past decade, biodegradable polymers or oligopeptides recognized by cell-surface receptors have been shown to increase drug specificity, lowering systemic drug toxicity in contrast to small-size fast-acting drugs. (Bai *et al*, 2008). The simplest calculations can be performed by hand, but computers are inevitably required to perform molecular modeling of any reasonably sized system. Molecular modeling and related computational techniques (often referred to as *in silico* approaches) are an integral part of the drug discovery and development workflow. Modeling approaches complement the development pipeline at a number of stages, but perhaps most significantly in the early phases of lead discovery and hit identification. The main objective is generally to discard the molecules less likely to be successful, hence focusing experimental work on the more valuable compounds. For this reason, *in silico* approaches are claimed to have a highly positive economic impact on the drug discovery process (3-4). Molecular modeling projects in this area will generally involve use of molecular modeling software to design inhibitor binding models. These models generate hit compounds, which are generally tested in relevant bioassays. Chemical modification will then usually be undertaken, with organic chemistry, biophysical measurements and molecular biology necessary.

The complexity of cancer and the vast amount of experimental data available have made computer-aided approaches necessary. Biomolecular modeling techniques are becoming increasingly easier to use, whereas hardware and software are becoming better and cheaper. Cross-talk between theoretical and experimental scientists dealing with cancer-research from a molecular approach, however, is still uncommon. This is in contrast to other fields, such as amyloid-related diseases, where molecular modeling studies are widely acknowledged.

Homology model of WNT16 B isoform cannot be found in public databases. Creating the 3D homology model of human WNT16B, we used Swissmodel (Waterhouse *et al*, 2018; Guex N *et al*, 2009; Benkert P, 2011; Bertoni M, 2017). As a template homology model *Helobdella sp*. WNT16B (Cho S *et al*, 2010) was used, to transform the known sequence of human WNT16B (McWhirte *et al*, 1999) into a model. Without the homology model, it is difficult to create docking scripts, because the sequence has not got the conformation information, which is essential to those *in silico* docking methods, where the programming language of the docking procedure is ideal for performing actions on large text types of datasets (like C, Perl, Python etc.). The analogy between human and *Helobdella sp*. WNT16B sequence is above 90 percent, so statistically the possible differences of the two homology model is under 10 percent, which is enough low, to use the mentioned *Helobdella sp*. WNT16B model, and to predict the usable human WNT16B model. The model was built using ProMod3 Version 1.1.0., in case loop modeling with ProMod3 fails, an alternative model is built using PROMOD-II (Guex N *et al*, 2009), the best result was found by BLAST (Camacho C *et al*, 2009). The model is a monomer polypeptide, with a CYS ligand added. The modeling was done in 2015 and in 2018, both effected the same result.

The docking methodology was done by Hex 8.0 spherical polar Fourier protein docking algorithm (Ritchie DW, 2005; Ritchie DW & Kemp GJL, 1999), the final sampling was calibrated to N=30, which effected the best result. All ligand interactions, bond-monitoring, and imaging were done using Discovery Studio 4.3.1 (BIOVIA DS, 2015). Hex was written mostly in C, but also uses some C++ for the GUI, the Discovery Studio uses Perl scripts. Results of binding interactions were inserted into XML tables, which can be easily used as database tables in SQL Server 2012 (Microsoft, 2012), with the help of SQL Server Management Studio 2016 (Microsoft, 2016).

*In vitro*, the tumor cell cultures were obtained from American Type Culture Collection and cultured with RPM1 medium 1640 or Leibovitz’s L-15 medium supplemented with 10% fetal calf serum. The cells were kept at 370C in a humidified atmosphere of 5% CO2 and 95% air.

*In vivo*, the tumors were implanted s.c. or were transplanted to the appropriate organs in the case of human tumors. In case of orthotopic tumor the tumor models were produced by transplanting cancer cells into an anatomically appropriate location in the host organism (Chou *et al*, 2010). Approximately 350,000 cells were cultured in 2 ml of culture medium in a well (6-well plates were used, Greiner, Nurtinger, Germany) and incubated for 24 h with different doses of AKOS004122375 in the absence of fetal calf serum. After trypsinization and dispersion, the cells were dyed with trypan blue and counted in a hemocytometer. The therapeutic efficacy of 4-(Furan-2-yl)hepta-1,6-dien-4-ol (AKOS004122375) in different long-term administration routes (once or twice a day for 2 weeks), the traditional injections (i.p., s.c.) versus infusion treatment (using an implantable Alzet type minipump), will be investigated later. Histochemical analysis of the different tumors from treated and control group animals were comparatively carried out.

During our toxicology studies, the effect of different doses of AKOS004122375 on survival and changes of body weight were tested for 2 weeks. The weight of different organs (heart, kidney, spleen, lung, liver, thymus and uterus) was measured after 1- and 18-day treatment, respectively). Hematological studies, the analysis of bone marrowth and cytology studies of different organs have been conducted as well.

Clonogenity assay provides a method to determine the surviving and colony forming ability of 4-(Furan-2-yl)hepta-1,6-dien-4-ol (AKOS004122375) treated adherent cell cultures. One thousand cancer or normal cells, at 70% confluence, will be seeded on 6-well plates in the required with 10% FBS. Cells were treated with the selected agents at IC90 concentration. After 3 days incubation the medium was changed a drug free one and surviving cell will cultured for 14-20 days. Cells were stained and fixed than the number of colonies, the number of cells in each colony and the morphology of cells was determined by confocal microscopy. Each experiment was performed two times: results will be accepted under 10% standard deviation. Results will be presented in surviving fraction percentage (SF), SF = No. of colonies after treatment/No. of colonies in untreated control x 100. SF values of tested agents on each cell line, complemented with the size of colonies and the morphology of cells. Tumor selectivity of tested agent can be determined with the help of a clonogenity assay on non-tumor (normal) cell lines.

The cytotoxicity of the compound AKOS004122375 was tested by reactions developed by our group consisting in the following strategies: end-point measurement and real-time evaluation of cell densities. In the first approach, we applied electronic pulse area analysis which has a high ability to detect the number of cells, to perform a cell size analysis and it provided the possibility to analyzing viability changes by the aid of altered permeability and electric moieties of the surface membrane. Long-term (24-72 h) types of impedance-based analysis are available not only for the characterization of chemotaxis and chemokinesis, but also it is a dedicated tool to follow changes in cell densities in the test chambers. In this case we can distinguish even fluctuation of cell numbers and characterize the cell cycle with high confidence.

The body weight of animals was measured and the tumor dimensions were measured with a microcaliper every second or third day. The tumor volume was calculated using the following formula: V= (×/6) x Lx D2 (V=Tumor volume, L: longest diameter, D: diameter perpendicular to L). Survival times related to that of the controls were recorded. Tumor volume measurements were continued until the first death in the control group. Mean values and standard deviations (S.D.) were calculated. Experimental data were subjected to computerized statistical analysis of variance with the Student-Newman-Keuls test; statistical significance was accepted at P>0.05 levels.

Apoptosis consists of a cascade of events lead to the ordered dismantling of critical cell survival components and pathways. Apoptosis events will be studied using fluorochrome conjugates of annexin-V to monitor changes in the cell membrane phospholipid asymmetry, thereby providing a convenient tool for detection of apoptosis cells. A distinctive feature of the early stages of apoptosis is the activation of cascade enzymes. These enzymes participate in a series of reactions that are triggered in response to pro-apoptotic signals and that result in the leverage of protein substrates, causes the disassembly of the cell. Caspase enzyme substrates and apoptosis kits can be used in these studies.

The National Institute of Oncology has a circle-like facility, personal background and infrastructure that are necessary complete the research project. The activity and research interest of the Department of Experimental Pharmacology of the National Institute of Oncology, Budapest, Hungary has been related to the following topics: breeding of laboratory animals (about 10 thousand mice per year, 60% of them are used in preclinical experiments), maintenance of different and transplantable mouse- and human tumor models (“tumor bank”), and preclinical examinations (e.g. toxicological, histological and hematological and different side effect examinations) of new antitumor agents and biological modulators. In addition to the Pharmacology Department, the object (2 modern laboratory and 3 experimental room) and personal (experienced researchers and support staff), conditions they are necessary for the conduct of experiments. For many years our department participates in different projects.

Specified pathogen-free mice (SPF) breeding of the Department of Experimental Pharmacology, National Institute of Oncology, Budapest, Hungary weighing 22-24 g used for these experiments. The animals were fed a sterilized standard diet (Biofarm) and had free access to tap water ad libitum. They were kept in macron cages at 23 −250 C (40-50% humidity), a lighting regiment of 12/12 hrs light/dark. The animals of this project were cared for according to the “Guideing Principles for the Care and Use of Animals” based upon the Helsinki declaration and were approved by the local ethical committee. In our experiments we used 7-10 mice/group (treatment and control groups).

## Acknowledgements

Our research was prepared with support from OTKA grant Hungary (grant # 115473), National Research, Development and Innovation Office Hungary, and National Institute of Oncology, Hungary.

## Author contributions

The authors have no interest in the referenced compounds vendors, computational program owners at all.

## Structured Methods

Reagents and Tools Table

| Reagent/Resource | Reference or Source | Identifier or Catalog Number |
| --- | --- | --- |
| **Experimental Models**<br>*List cell lines, model organism strains, patient samples, isolated cell types etc. Indicate the species when appropriate.* |  |  |
| In vitro cell colonies | National Institute of Oncology, Hungary | obtained from American Type Culture Collection, cultured with RPM1 medium 1640 or Leibovitz's L-15 medium supplemented with 10% fetal calf serum |
| Specified pathogen free mice (SPF) | National Institute of Oncology, Hungary | weighing 22-24 g |
| **Oligonucleotides and other sequence-based reagents**<br>*For long lists of oligos or other sequences please refer to the relevant Table(s) or EV Table(s)* |  |  |
| 2JWA<br>ErbB2 transmembrane segment dimer spatial structure, Homo Sapiens (Human), HER2 TMD homodimer | Bocharov et al, 2008 | pdb ID: 2JWA<br>PubMed: 18178548<br>pdb file |
| 2M0B Homodimeric transmembrane domain of the receptor tyrosine kinase ErbB1 (EGFR, HER1) in micelles, Homo Sapiens (Human), HER TMD homodimer | Bocharov et al, 2016 | pdb ID: 2M0B<br>PubMed: 26903218<br>pdb file |
| WNT16B Homo Sapiens (Human) sequence | McWhirter et al,1999; | UniProtKB - Q9UBV4 (WNT16_HUMAN), sequence |
| WNT16B Homo Sapiens (Human) pdb | This study | pdb file |
| WNT16B Homo Sapiens (Human) docking into the active form of ErbB2 transmembrane segment dimer spatial structure Homo Sapiens (Human) 2JWA | This study | pdb file |
| 4-(Furan-2-yl)hepta-1,6-dien-4-ol (AKOS004122375) docking into the mutant complex of ErbB2 transmembrane segment dimer spatial structure Homo Sapiens (Human) 2JWA and WNT16B Homo Sapiens (Human) | This study | pdb file |
| 4-(Furan-2-yl)hepta-1,6-dien-4-ol (AKOS004122375) docking into ErbB2 transmembrane segment dimer spatial structure Homo Sapiens (Human) 2JWA | This study | pdb file |
| WNT16B Helobdella sp. SJC-2009 sequence | Cho SJ et al, 2010 | UniProtKB - D6N3F7 (D6N3F7_9ANNE), sequence |
| WNT16B Homo Sapiens (Human) docking into the homodimeric transmembrane domain of the receptor tyrosine kinase ErbB1 (EGFR, HER1) in micelles, Homo Sapiens (Human) 2M0B | This study | pdb file |
| 4-(Furan-2-yl)hepta-1,6-dien-4-ol (AKOS004122375) docking into the homodimeric transmembrane domain of the receptor tyrosine kinase ErbB1 (EGFR, HER1) in micelles, Homo Sapiens (Human) 2M0B | This study | pdb file |
| 4-(Furan-2-yl)hepta-1,6-dien-4-ol (AKOS004122375) docking into the mutant complex of the homodimeric transmembrane domain of the receptor tyrosine kinase ErbB1 (EGFR, HER1) in micelles, Homo Sapiens (Human) 2M0B and WNT16B Homo Sapiens (Human) | This study | pdb file |
| **Chemicals, Enzymes and other reagents**<br>(e.g. drugs, peptides, recombinant |  |  |
| *proteins, dyes etc.)* |  |  |
| 4-(Furan-2-yl)hepta-1,6-dien-4-ol | AKos Consulting & Solutions GmbH | AKos Consulting & Solutions ID: AKOS004122375<br>PubChem CID: 108669579 |
| **Software**<br>*Include version where applicable* |  |  |
| Discovery Studio 4.3.1. | BIOVIA DS, 2015 | Release 4 |
| Hex 8.0. | Ritchie DW, 2005; Ritchie DW & Kemp GJL, 1999 | Release 8 |
| SQL Server 2012, Transact-SQL (T-SQL) | Microsoft | Version 2012 |
| SSMS 2016, SQL Server Management Studio | Microsoft | Version 2016 |

